# Quantitative, temporal dynamics of the cellular amino acid economy reveals public vs private goods and enables syntrophic yeast community design

**DOI:** 10.1101/2025.07.09.664033

**Authors:** Shabbir Ahmad, Sunil Laxman

## Abstract

The biosynthetic capacity of the cell governs the production and exchange of amino acids. Amino acids have diverse metabolic origins and intracellular requirements. Therefore, to understand the cellular amino acid economy we require a quantitative understanding of intracellular and extracellular their amounts, and how distinct amino acids change temporally during growth. Using the unicellular eukaryote, yeast, we establish a quantitative blueprint of the intracellular and extracellular amino acid economy, including the fluxes of production, secretion and consumption across different growth phases. Certain amino acids dominate the intracellular pool, and proportions of distinct amino acids continuously change during growth. The extracellular pool is notably distinct from the intracellular pool in terms of composition and dynamics. Only some public amino acids are subsequently consumed, while others are secreted in surplus of cellular demands. Five amino acids are intracellular and privatized, and continuously utilized to support diverse metabolism. Through this, we establish pairs of highly effective synthetic, exchange-based communities of public good auxotrophs. Our results suggest organizing frameworks addressing the dynamic intracellular and inter-cellular amino acid economy and its trade, and informs the rational engineering of syntrophic cell communities.

## Introduction

Cells make, utilize and accumulate metabolic resources such as amino acids in substantial amounts, and these resources can also be secreted outside to be traded between cells. Enabled by such trade, cells assemble into syntrophic communities – as seen in diverse microbial systems (1–3), or between cells in tissues. The consequences of metabolic exchange between cells include distributed metabolic functions between cell groups, divided metabolic labor, increased metabolic productivity, or reduced biosynthetic load (1, 3–5). This suggests that by rationally manipulating the exchange of amino acids, it may be possible to design engineered cells that assemble and form exchange-based consortia, or can support complex biosynthesis. Through division of labor, cells create metabolic efficiencies between each other. This is commonly observed in microbial communities, which host auxotrophic cells that rely on metabolites produced by other cells, creating exchange based syntrophic consortia (3, 6–13). A spectacular extreme example in nature are aphid insect-bacterial endosymbionts, who survive based on extensive amino acid trade and associated metabolic optimizations(14, 15). Metabolic trade also occurs in colonies of prototrophic cells (which can biosynthesize their own metabolites), through the self-organization of cells into groups with division of labor to share resources (1, 16–19). Thus, building metabolic efficiencies based on biochemical and thermodynamic constraints can drive the assembly of cell communities (3, 19, 20). However, at a more fundamental level, the biochemical constraints that determine which metabolic resources are exchanged between cells, and how much cells prioritize major resources such as amino acids remain poorly understood.

Amino acids are very commonly exchanged between cells (5, 21). But amino acids are themselves not homogeneous entities. Instead, they are chemically, metabolically and functionally diverse, have varying individual cellular requirements, and are present in distinct intracellular amounts (22, 23). Cells also maintain different extents of strategic amino acid reserves for metabolic and translational requirements (24). Indeed, cells function as demand-driven economies with respect to different amino acids – and the net demand for each amino acid shapes hierarchies or priorities of amino acid resource allocations (22, 25). In order for a resource to be traded, sufficient quantities need to be available in excess of demand. The amounts of these amino acids will inherently be a function of their cycles of production – consumption – secretion. Therefore, in order to build a first-principles basis of understanding the cellular amino acid economy and trade, we require quantitative estimates of the amounts of intracellular amino acids in the context of extracellular pools, and how these pools change temporally across the growth phases of a cell. This knowledge can also aid in designing biosynthetic processes in engineered cells to produce amino acid derived molecules. However, there is limited quantitative information available for temporal changes in amino acid production, consumption and secretion across different phases of cell growth, even for prototrophic cells that make their own amino acids from carbon and nitrogen precursors.

In this study, we utilized the eukaryotic model cell *Saccharomyces cerevisiae*, which is prototrophic for the biosynthesis of all amino acids, to quantitatively determine intracellular and extracellular amino acid amounts across a 24-hour growth phase. Estimates of biosynthetic flux, along with absolute quantification of amino acids revealed that cells maintain distinct amounts and magnitudes of intracellular and extracellular amino acids that do not correlate with each other in abundance, demand or hierarchies of utilization. Specific amino acids are strictly privatized and not secreted to the external medium, and these privatized amino acids are continuously required to support critical intracellular metabolic processes. A group of amino acids are secreted as public goods into the extracellular environment, only some of which are subsequently taken up and utilized to support metabolism of the producing cell. This quantitative blueprint of public and private good amino acids enabled the rational design of paired communities of public good amino acid auxotrophs, based on the amounts of amino acids exchanged. Pairs of auxotrophs of surplus public good amino acids established stable, syntrophic communities that grow robustly together. Our data therefore reveal quantitative hierarchies of cellular amino acid production, accumulation and secretion, with implications for the design of synthetic consortia for metabolic engineering applications.

## Results

### High amino acid production is limited to the early phase of exponential growth

The intracellular amounts of amino acids reflect the net amount of production, versus consumption and secretion of amino acids. Understanding this biosynthesis– secretion – consumption cycle across cell growth, and as cells enter stationary phase is a fundamental question in quantitative cellular physiology (Fig. 1A). However, even in laboratory based model microbial culture settings, we have a limited understanding of amino acid production-secretion-consumption amounts and temporal changes therein (Fig 1A). We therefore addressed this question, using prototrophic yeast cells grown in defined minimal medium without supplemented amino acids. In this condition, prototrophic yeast carry out *de-novo* biosynthesis of all amino acids, thereby allowing a quantitative assessment of amino acids over time, across phases of growth. In this study, we use a robust, prototrophic yeast strain (CEN.PK background), growing in batch cultures in synthetic, minimal medium with 110 mM glucose and ∼35 mM ammonium sulfate as sole carbon and nitrogen sources respectively in all experiments.

**Fig 1.**
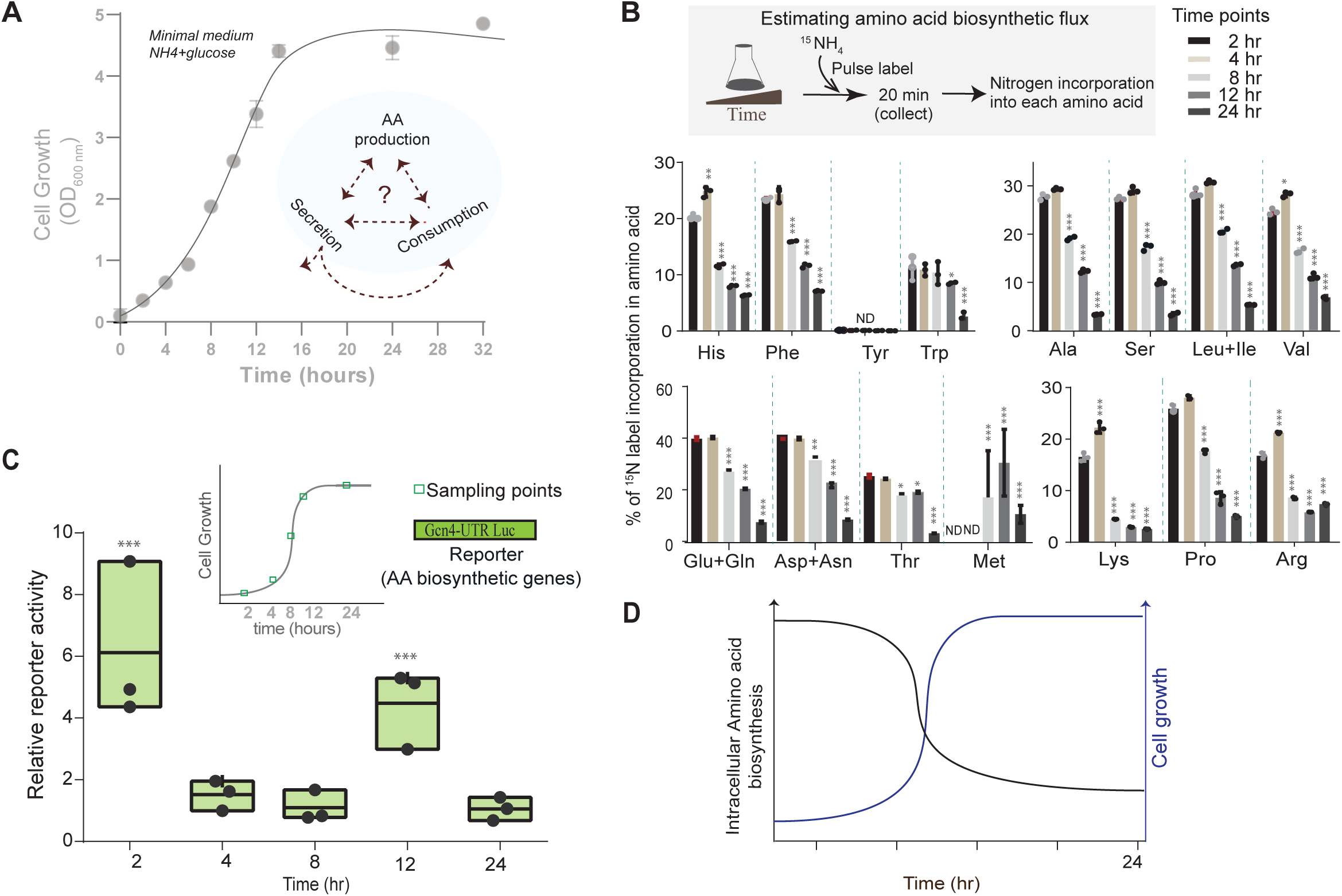
High amino acid production is limited to the early phases of cell growth. **(A)** Cells modulate and balance amino acid production (and acquisition), secretion and consumption during different growth phases, but the dynamics and interplay of this production-secretion-consumption is poorly understood. On the right, the schematic shows a standard growth curve of *S. cerevisiae* in batch culture over ∼24 hours (in synthetic, defined, minimal medium with glucose and ammonium sulfate as the sole carbon and nitrogen sources), and also indicates time points at which samples were collected, to assess intracellular and extracellular amino acid pools. **(B)** Estimates of relative amino acid biosynthetic flux across distinct growth phases: (Upper inset) - Experimental workflow used to estimate amino acid biosynthetic flux, using a pulse-label of ^15^N labelled ammonium sulfate, and measuring label incorporation into the respective amino acid using targeted LC-MS/MS, from cells collected at the indicated time points. The relative label incorporation into each newly synthesized amino acid is shown. The amino acids are presented as groups loosely based on their metabolic origins. Data are reported showing three biological replicates. Comparisons are to the 2hr time point of each sample. Significance by Student’s t-test. * p<0.05, ** p<0.01, *** p<0.001. ND – not detected. **(C)** Schematic summarizing the relative amino acid biosynthetic flux in cells versus biomass formation over time, where cells have high biosynthetic flux for all amino acids during early/exponential growth, and very low biosynthesis during later growth phases. **(D)** Global transcriptional induction of amino acid biosynthetic genes across growth phases: The extent of transcriptional activation of amino acid biosynthetic genes in distinct phases of growth was estimated using the activity of an established transcriptional reporter (Gcn4-luc)(Gupta et al., 2019). Cells were grown in synthetic minimal medium, as indicated in panel 1A. Normalized relative reporter activity is reported (n=3, mean ± SD).

Our first step was to quantify the changes in amino acid biosynthetic flux during different phases of cell growth. For this, we designed experiments to estimate amino acid biosynthetic flux over a 24-hour period of cell growth (Fig. 1B). To quantitatively assess the relative rate of amino acid biosynthetic flux over this 24-hour window, we designed a metabolic flux experiment, where employed a short pulse of stable-isotope labelled nitrogen precursors at the indicated time point, followed by assessing the relative nitrogen label incorporation into newly synthesized amino acids (Fig. 1B). For this, ^15^N-labeled ammonium sulfate was pulsed briefly into cells (at the indicated time points), cells were collected, and the incorporation of ^15^N into newly synthesized amino acids was estimated by quantitative, targeted LC-MS/MS. This allowed us to quantify and compare relative amino acid biosynthetic fluxes at different time points, viz. 2, 4, 8, 12, and 24 hours (Fig. 1B). We observed highest amino acid biosynthetic flux during the first 2-4 hours of cell growth (Fig. 1B), which declines substantially after ∼12 hours of growth (Fig. 1B. These data establish that cells in logarithmic phase of growth have highest amino acid biosynthesis.

We also assessed how this actual amino acid biosynthetic flux was reflected at the level of amino acid biosynthetic gene expression. The expression of amino acid biosynthetic genes are commonly used as a proxy to indicate amino acid biosynthesis, although this cannot distinguish reduced substrate availability, or starvation based feedback responses where amino acid biosynthetic genes can increase. To assess amino acid biosynthetic gene expression, we used a well-established luciferase based reporter for amino acid biosynthesis (22, 25), based on the activity of the amino acid master-regulating transcription factor Gcn4 (25–28) (Fig. 1C). This reporter activity was measured at 2, 4, 8, 12, and 24 hours, and it revealed two distinct waves of expression of amino acid biosynthetic genes (Fig. 1C). Reporter activity was highest during the early growth phase (2 hours), decreased thereafter, and a second burst of activity followed post ∼12 hours (Fig 1C). However, as observed earlier (Fig. 1B), amino acid biosynthetic flux is high only in the early (2-4 hr) periods of growth, and is low at ∼12 hours. Therefore, the second wave of transcription would be consistent with a starvation response, where cells sense reductions in actual amino acid biosynthetic flux as nutrients deplete, and upregulate transcriptional programs for amino acid biosynthetic genes.

Collectively, these data comprehensively establish the temporal dynamics of amino acid production, highlighting that the majority of biosynthesis is restricted to early phases of growth.

### Amounts and hierarchies of intracellular and extracellular amino acids are distinct and uncorrelated

Given the temporal restriction of amino acid biosynthesis, we next asked what are the intracellular and extracellular amino acid concentrations during different growth phases. This is an important, quantitative question asking how much the amounts differ for distinct amino acids. Thereby, this can address what the intracellular demand versus production of different amino acids are, and identify which amino acids might be shared in the external environment. We used quantitative, targeted LC-MS/MS approaches to determine absolute intracellular amino acid amounts across these (2, 4, 8, 12, and 24 hour) phases of growth, and estimated the per cell concentration based on cell number and volume (Fig. 2A, and Table 1) (also see materials and methods for calculations). Additionally, the range of intracellular amino acid amounts of different amino acids at 4 hours of growth (when cells are in log-phase growth) are illustrated as a heat map (Fig. 2B), highlighting the large, order-of-magnitude differences in amounts of different amino acids. The absolute intracellular amino acid concentrations differed for the most abundant to least abundant amino acids by ∼four orders of magnitude (Fig. 2A). Consistent with studies from bacteria (23), and recent predictions (22) the most abundant amino acids were glutamate, glutamine, alanine and aspartic acid, while methionine was among the lowest across different phases of growth (Fig. 2A, 2B, Table 1). Note: cysteine was not quantified across these times due to its redox sensitivity. However in similar media conditions cysteine concentration are in the high nM range, lower than methionine (29). Interestingly, over this temporal window the absolute intracellular concentrations of amino acids, as well as relative proportions of each amino acid as a fraction of total amino acids inside a cell change markedly (Fig. 2C, Table 1). Glutamate and glutamine remain the most abundant intracellular amino acids across this time-window, but their amounts decrease steadily. Some amino acids, notably aspartate, exhibited a decline across the growth phase (Fig. 2A, 2C, Table 1). Certain other amino acids, such as histidine, phenylalanine, tyrosine, asparagine, lysine, and the branched chain amino acids, accumulated throughout growth (Fig. 2C), while amino acids like proline initially accumulated but later decreased. Interestingly, intracellular arginine levels were maintained over time (Fig. 2A, 2C, Table 1). A few amino acids, particularly phenylalanine, tyrosine, tryptophan, valine, leucine/isoleucine, and asparagine substantially increased after 12 hours of growth (Fig. 2C, Fig. S1).

**Fig. 2:**
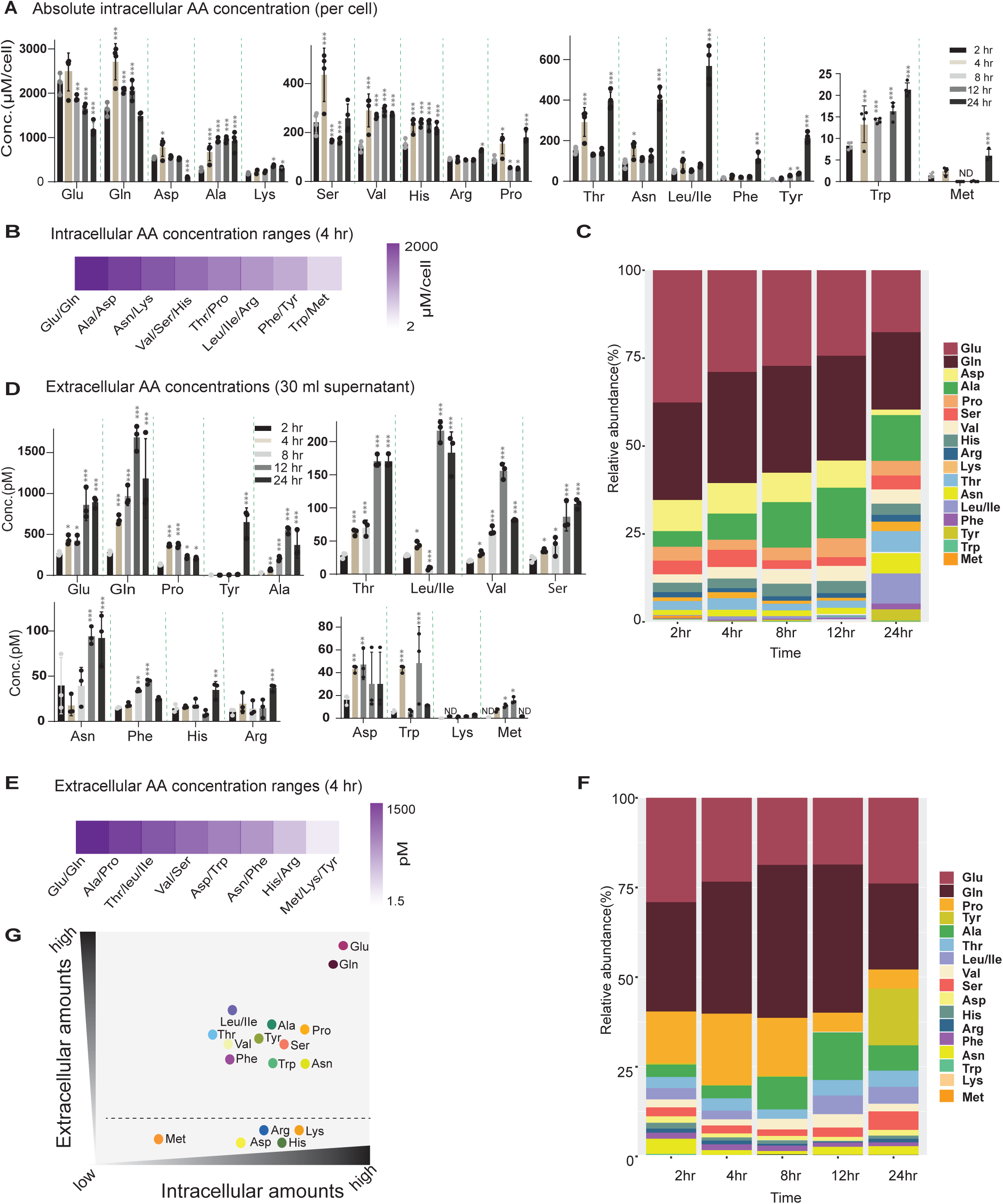
Amounts and hierarchies of intracellular and extracellular amino acids are distinct and uncorrelated. A. Absolute concentrations of intracellular amino acids from cells collected at the indicated growth phases. Intracellular amino acids were quantitatively estimated by LC-MS/MS. The amino acids are arranged based on the order of their intracellular concentration (high/intermediate/low in (μM/cell). The estimated volume of a single yeast cell is 40fl. N=4 (biological replicates). Significance comparisons are to the 2hr time point of each sample (Student’s t-test). * p<0.05, ** p<0.01, *** p<0.001. B. A heatmap to illustrate relative abundances of groups of amino acids inside a cell after four hours of growth. The purple color intensity reflects the amount of amino acid present, over three orders of magnitude (μM/cell). C. Proportions of intracellular amino acids over time at different cell growth phases: Stacked bar plots illustrate changes in the relative proportions of distinct intracellular amino acids across various phases of growth. Note that the relative proportions of several amino acids change substantially at different times. Also see Fig. S1. D. Absolute concentrations of extracellular amino acids (accumulated in the medium) across the indicated times of growth. Measurements were made in 30 ml of culture medium volume. The extracellular amino acids are arranged from the most to least abundant at the indicated time points. Note that while glutamate and glutamine remain the most abundant extracellular amino acids, several other amino acids in the extracellular environment are present at amounts that do not correlate with their intracellular amounts (panel 1B). Some amino acids are not detectable or present at very small amounts in the extracellular environment. N=3, mean ± SD (biological replicates). Significance comparisons are to the 2hr time point of each sample (Student’s t-test). * p<0.05, ** p<0.01, *** p<0.001. E. A heatmap to illustrate relative abundances of groups of amino acids outside a cell in the extracellular medium after four hours of growth. The purple color intensity reflects the amount of amino acid present, over three orders of magnitude. The concentrations are in pM present in 30 ml of culture medium. F. Proportions of extracellular amino acids over time across growth phases: Stacked bar plots illustrate changes in the relative proportions of distinct intracellular amino acids across various phases of growth. Note that the relative proportions of several amino acids change substantially at different times. Also see Fig. S2. G. A schematic illustration showing relative amounts and proportions of intracellular and extracellular amino acids, observed after twelve hours of cell growth in minimal medium. Many amino acids are present in abundance in both the intracellular and extracellular environments, while some others (Asp, Met, Arg, Lys, His) are present in distinct concentrations inside a cell, but are almost entirely absent in the extracellular environment.

**Table 1:** Absolute intracellular and extracellular amino acid concentration over time.

| **Intracellular amino acid concentrations (µM)** |  |  |  |  |  |
| --- | --- | --- | --- | --- | --- |
| **Amino acid** | **2 hours** | **4 hours** | **8 hours** | **12 hours** | **24 hours** |
| Glu | 2229 | 2479 | 1882 | 1650 | 1210 |
| Gln | 1636 | 2709 | 2072 | 2058 | 1504 |
| Asp | 546 | 779 | 559 | 518 | 105 |
| Ala | 268 | 671 | 930 | 954 | 894 |
| Lys | 192 | 246 | 234 | 368 | 284 |
| Ser | 241 | 436 | 166 | 167 | 271 |
| Val | 142 | 290 | 266 | 290 | 274 |
| His | 152 | 232 | 239 | 235 | 34 |
| Arg | 91 | 86 | 87 | 88 | 131 |
| Pro | 89 | 154 | 56 | 52 | 184 |
| Thr | 151 | 291 | 130 | 142 | 405 |
| Asn | 88 | 162 | 110 | 123 | 418 |
| Leu/Ile | 48 | 89 | 51 | 78 | 589 |
| Phe | 15 | 24 | 18 | 21 | 40 |
| Tyr | 8 | 13 | 28. | 38 | 218 |
| Trp | 8 | 13 | 14 | 16 | 21 |
| Met | 1 | 2 | ND | ND | 5 |
| **Extracellular amino acid concentration (pM) (from 30 ml culture volume)** |  |  |  |  |  |
| **Amino acid** | **2 hours** | **4 hours** | **8 hours** | **12 hours** | **24 hours** |
| Glu | 245 | 416 | 391 | 710 | 899 |
| Gln | 254 | 663 | 905 | 1502 | 915 |
| pro | 128 | 378 | 335 | 203 | 206 |
| Tyr | 3 | 3 | 5 | 9 | 734 |
| Ala | 29 | 58 | 175 | 501 | 256 |
| Thr | 23 | 65 | 74 | 164 | 165 |
| Leu/Ile | 27 | 43 | 9 | 198 | 204 |
| Val | 19 | 30 | 61 | 142 | 80 |
| Ser | 26 | 33 | 46 | 73 | 112 |
| Asn | 28 | 30 | 16 | 85 | 60 |
| Phe | 15 | 16 | 32 | 40 | 38 |
| His | 14 | 15 | 16 | 12 | 39 |
| Arg | 12 | 15 | 10 | 3 | 39 |
| Asp | 18 | 45 | 7 | 12 | 15 |
| Trp | 7 | 3 | 11 | 18 | 11 |
| Lys | ND | ND | ND | ND | 3 |
| Met | 1 | 1 | 1 | 1 | 1 |

These results suggest that (i) many amino acids are produced and secreted in amounts exceeding cellular demand, and (ii) the relative or proportional pools of distinct amino acids within a cell are not constant, but vary depending on the growth state of the cell. Therefore, this immediately raises the question on the nature of the extracellular amino acid environment across this temporal window, in order to understand which amino acids are secreted and consumed by cells. We therefore quantified absolute amounts of extracellular amino acids from the culture supernatant (measured in picomolar amounts in 30 mL of liquid culture), across growth phases. Amino acids were arranged in groups from the most to the least abundant after 4 hours of growth (Fig. 2D, Table 1). Interestingly, only a group of amino acids were present at significant amounts in the extracellular medium (Fig. 2D). Similar to intracellular trends, glutamine and glutamate were the most abundant extracellular amino acids, followed by alanine. Further, proline, branched chain amino acids, and threonine were present in substantial amounts in the extracellular medium (Fig. 2D, 2E, Table 1). In addition, serine, asparagine, threonine, and leucine/isoleucine showed a consistent increase in the extracellular environment (Fig. 2D), also consistent with their intracellular accumulation patterns (Fig. S1). Cells therefore secrete significant amounts of select amino acids during logarithmic growth, when nutrient availability is high (Fig. 2D, Fig. 2E). Additionally, histidine, arginine, phenylalanine, and serine displayed minor variations in their extracellular concentrations, while having considerable intracellular concentrations (Fig. 2F; S2).

In contrast however, some amino acids, particularly histidine, arginine, aspartic acid, methionine, and lysine were present in very low or undetectable amounts in the extracellular medium (Fig. 2D, 2E & S2, Table 1). These results underscore that the extracellular amino acid environment is not homogeneous throughout growth (Fig. 2F). Furthermore, over time the levels of a subset of extracellular amino acids decrease, and the relative proportion of extracellular amino acids changed accordingly (Fig. 2F). This suggests phases of subsequent reuptake of these select amino acids (Fig. 2F, Fig. S2). Two important conclusions can be drawn. First, the extracellular environment does not reflect the intracellular amino acid environment – either in terms of the absolute amounts of distinct amino acids, or in terms of the relative proportion of availability of each amino acid. Importantly, some amino acids, which may be fairly abundant inside the cell, are almost unavailable outside. Second, note that while many of the abundantly secreted amino acids have high biosynthetic costs (proline, the branched chain amino acids, and threonine), they all have fairly modest demand within growing cells, as extensively shown earlier (22). This is further discussed later in the manuscript. This reiterates that demand-driven criteria drive the cellular amino acid economy, and prioritizations (22).

We synthesize this qualitative relationship between intracellular and extracellular amino acids, and visually illustrate this in Fig. 2G (reflecting twelve hours of growth). From this, we can observe that only a few amino acids show similar patterns in their intracellular and extracellular amounts, while others remain uncorrelated (Fig. 2G). Of note, arginine, lysine, histidine, and aspartic acid are abundant intracellular amino acids, but are low/undetectable in the extracellular environment, revealing that these amino acids are not secreted in substantial amounts (Fig. 2G). In contrast, a subset of amino acids –phenylalanine, tryptophan, tyrosine, leucine/isoleucine, proline, threonine and asparagine substantially accumulate in the extracellular environment over time. Note that all these amino acids have varying costs of synthesis (22), and no association can therefore be made with the costs incurred in making these molecules. However, all of these amino acids have relatively low cellular demand, and production can therefore exceed cellular requirements (22). Collectively, these results reveal how the dynamic extracellular amino acid environment differs strikingly from the intracellular amino acid environment.

### Distinct amino acids function as a private, public or surplus goods

These data suggest that instead of forming homogeneous pools, the different amino acids can be divided broadly into two categories: (i) as a private good, and (ii) two types of public goods (Fig. 3A). In economic terms, public goods are resources that are available to all members of a population, while private goods are those that are exclusively used only by the producing individuals (Fig. 3A). For cells, this means that the amino acids that accumulate in the extracellular environment are public goods (Fig. 3A), while those that are restricted within the intracellular environment are privatized goods. A group of amino acids are present in low/undetectable amounts in the extracellular medium, despite being present at considerable intracellular amounts, indicating privatization of these metabolites (Fig. 3B). This set of privatized amino acids comprised of aspartic acid, lysine, arginine, histidine, and methionine (Fig. 3B). Contrastingly, the remaining amino acids fell into two main types of public goods – those that accumulate/do not decrease in the extracellular environment over time, and others that accumulate at earlier time points but subsequently deplete over time, thus suggesting possible uptake and utilization (Fig. 3B). Asparagine, threonine, leucine/isoleucine, tyrosine substantially increased in the extracellular medium (as well as intracellularly), indicating that these amino acids are public goods (surplus). Of the other amino acids, alanine and glutamine initially accumulate in the extracellular medium, but subsequently decline, suggesting later demand and utilization (Fig. 3B). These are classified as public goods (consumed). Similar trends (at lower concentrations) can be observed for phenylalanine, proline, tryptophan, glutamine, alanine and valine (Fig. 3B). Therefore, prototrophic yeast cells clearly treat distinct amino acids differently –as private goods, public good-surplus, and public good-consumed (Fig. 3B). The private or public nature of the amino acid can therefore be unambiguously defined by the extracellular amounts alone.

**Fig. 3:**
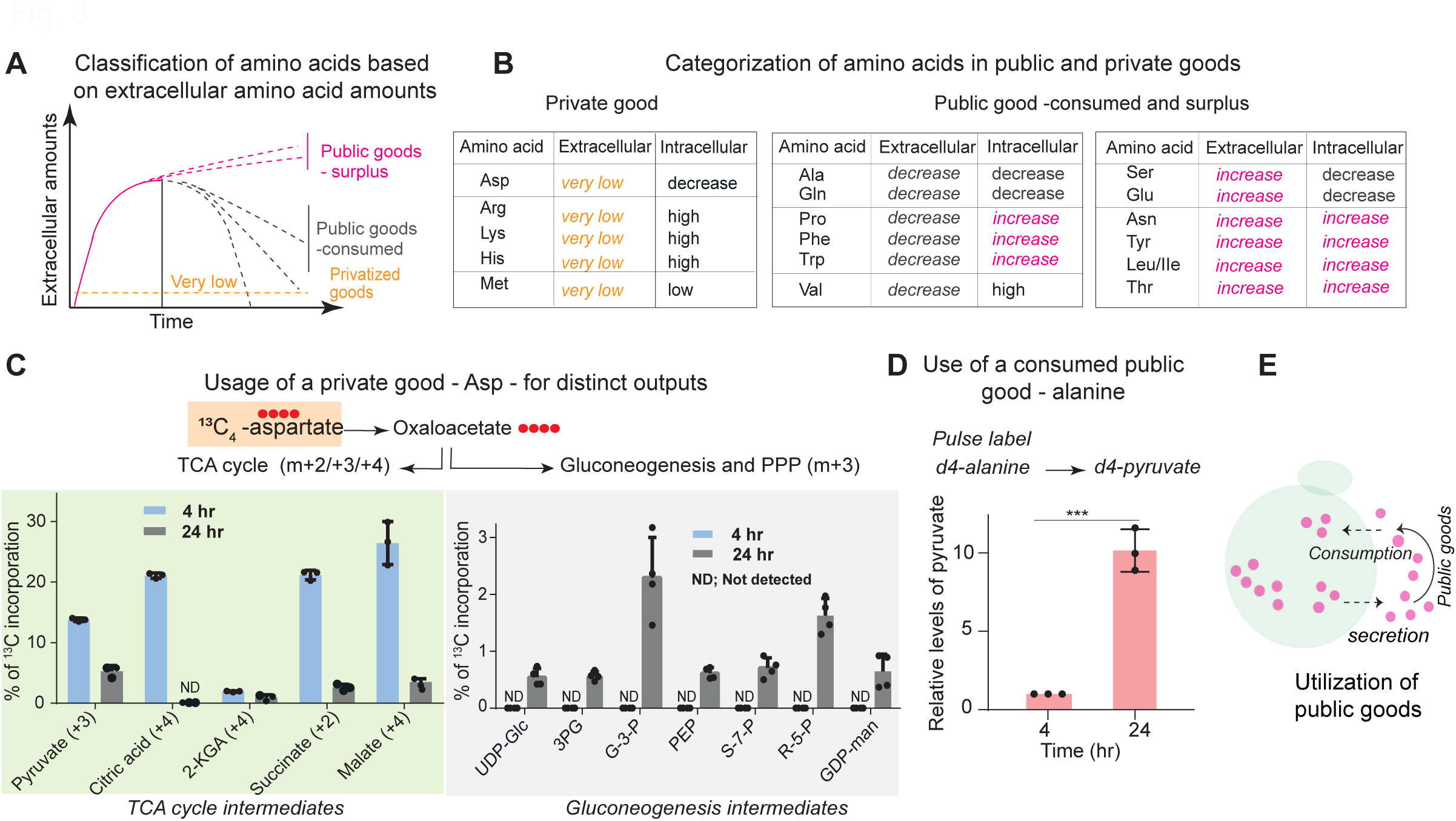
Distinct amino acids function as a private, public or surplus goods. (A) Classification of amino acids as private or public goods based on extracellular amounts and availability. Amino acids can be classified as private goods, or two types of public goods (surplus or consumed) based on extracellular availability over time. The measured extracellular pool of the respective amino acid is an indication of the balance of production and reuptake/consumption of the same. Privatized amino acids are not present in significant amounts in the extracellular culture medium. Surplus public goods will continuously accumulate in the extracellular culture medium over time, while consumed public goods will initially accumulate in the extracellular medium and subsequently decrease in abundance. (B) Categorization of individual amino acids as private or public goods. This grouping of amino acids is based extracellular availability, as well as changes in the extracellular amounts temporally. Private good amino acids include Asp, Met, His, Arg, Lys – and are shaded in light amber area. These are present in very low amounts in the extracellular medium. Amino acids that classify as public goods – consumed include Ala, Gln, Pro, Phe, Trp, Val. These first accumulate and then decrease over time in the extracellular medium. Public goods-surplus include Ser, Asp, Tyr, Leu/Ile, Thr. These increase/remain high in the extracellular medium. Also see Fig. S3 for privatized amino acids. (C) Demonstration of distinct allocations of a private good (aspartate) over time: Experimental design to track the metabolic fates of aspartate at different times, using a ^13^C_4_-aspartate pulse-label to monitoring incorporation into distinct branches of carbon metabolism – the TCA cycle or gluconeogenesis. Aspartate converts to oxaloacetate, and the carbon from aspartate subsequently can be tracked as it incorporates into TCA cycle or gluconeogenic intermediates by targeted LC-MS/MS. The relative utilization of aspartate as a carbon donor for intermediary metabolites in the TCA cycle or gluconeogenesis at two different times of cell growth was estimated. The grey shaded box is for gluconeogenesis intermediates, and the green shaded box is for the TCA cycle, comparing cells after 4hr or 24hr of batch culture growth. Note that in 4 hr, cells primarily use oxaloacetate from aspartate for the TCA intermediates, and after 24hr mainly use aspartate for gluconeogenesis. Data are from three biologically independent experiments (n=3), shown as mean ± SD. Also see Fig. S3. (D) Demonstration of the use of a public good (alanine) from the extracellular medium after 24 hours: Left inset – a schematic showing how a utilized public good will function. Right - Relative consumption of exogenously provided alanine via conversion to pyruvate. As shown in the experimental design (top), cells at 4hr and 24hr of growth were spiked with d_4_-alanine, and the relative d_4_- label conversion into pyruvate was estimated using targeted LC-MS/MS. Bar plot was shown relative levels of d_4_-pyruvate. The pyruvate label coming from alanine considerably increases after 24hr. Data are from three biologically independent experiments (n=3), shown as mean ± SD, *p<0.05, ***p<0.0001 (student’s t-test). Also see Fig. S3. (E) A schematic of cells described the amino acid, secretion and their uptake/exchange, when cells grown in the minimal medium.

We also assessed the biosynthetic costs of the privatized goods (Fig. S3), based on detailed estimates made earlier (22). However, the privatized amino acids did not associate with high costs –aspartic acid, arginine and lysine are amino acids with relatively low costs, methionine has an intermediate cost of synthesis, while histidine has high biosynthetic costs (22) (Fig. S3). In contrast, when assessed based on their cellular demand for multiple processes, the privatized amino acids all had continuous, changing, and high demands for multiple, distinct outputs as shown earlier (22), as also as illustrated in Fig. S3 which shows demand quantifications and usage respectively. Notably, all privatized amino acids have critical, continuous usage requirements in cells across stages of growth. Aspartate is an important, metabolically flexible precursor that fuels the tricarboxylic acid (TCA) cycle, and enables gluconeogenesis via its conversion to oxaloacetate (Fig. 3C, S3), and is essential for nucleotide synthesis (17), making it metabolically useful under diverse nutrient environments. Arginine has high, continuous demand for polyamine synthesis, and ribosome biogenesis, particularly during metabolic transitions (Fig. S3). Histidine is a precursor in the tetrahydrofolate cycle, plays a role in metal chelation, and also maintains cellular buffering (Fig. S3), and therefore is continuously required. Methionine is the precursor for S-adenosyl-methionine (SAM) and is continuously fluxed to SAM biosynthesis required for most biological methylation, SAH, polyamine synthesis, and the folate cycle (Fig. S3), making it a continuously utilized metabolite. Thus, the privatized amino acids are all continuously allocated towards multifaceted metabolic processes that the cell requires continuously (Fig. S3).

To experimentally assess the usage of privatized amino acids, we used aspartate as an example of a resource with diverse allocations at different times. For this, we carried out stable isotope-based label-incorporation experiments to estimate relative aspartate allocations during the course of growth, towards the TCA cycle or gluconeogenesis (Fig. 3C). We monitored label incorporation of ^13^C_4_-aspartate pulsed during early (4 hours) or late (24 hours) growth towards the TCA or gluconeogenesis at different phases of growth (Fig. 3C). At 4 hours, the labelled ^13^C_4_-aspartate was predominantly incorporated into the intermediates of the TCA cycle (Fig. 3D, light green box). This indicates that aspartate is continuously converted to oxaloacetate and supports TCA activity during the exponential growth phase. In contrast, at 24 hours, carbon from aspartate was incorporated substantially into gluconeogenesis intermediates (Fig. 3C, grey box) including UDP-Glc, PEP, G-3-P, GDP-Man, R-5-P, 3PG, and S-7-P (Fig. 3C). These data demonstrate the continuous, changing use of aspartate to sustain diverse cellular needs, consistent with a plausible requirement to privatize it.

A subset of publicly accumulated amino acids decreased in the extracellular environment after ∼24 hours (late growth). These suggest that cells excrete these as public goods, but might uptake and utilize them later, as the external environment changes. It is experimentally prohibitive to demonstrate this for each amino acid. However, we use alanine as an example to experimentally test this. Alanine accumulates in the extracellular medium at early time points, but start to decrease after 24 hours (Fig. 2D). Alanine is synthesized from pyruvate (from glucose metabolism), and alanine is catabolized to pyruvate, which can subsequently re-enter central carbon metabolism. To assess the use of alanine as an example of a late-utilized public good, we added a pulse of stable isotope labelled (d4-alanine) alanine, and estimated (uptake and utilization) relative flux towards pyruvate at 4 hours and 24 hours. Pyruvate formation from alanine was substantially higher at 24 hours compared to 4 hours, indicating robust uptake and utilization of alanine for central carbon metabolism after 24 hours of growth (Fig. 3D). These findings illustrate how secreted alanine functions as a utilized public good, to support central carbon metabolism during late growth phases.

Collectively, our data establish that prototrophic cells produce and secrete select amino acids in excess, as public goods (Fig. 3E). Some of these public goods are subsequently taken up and utilized or exchange by the cells, based on subsequent metabolic requirements. Contrastingly, tightly privatized amino acids are restricted to the intracellular environment, and can be continuously used for multiple, changing resource allocations.

### Cells secrete sufficient excess of amino acids to support growth of public good auxotrophs

Our data lead to the hypothesis that cells easily obtain public good amino acids from the external environment, and therefore auxotrophs of these amino acids would sustain their requirements exclusively from this external source (Fig. 4A). As a corollary, our data directly predicts that auxotrophs of private good amino acids would not sustain growth in spent medium (Fig. 4A). We therefore investigated if the secreted amino acids in spent media could sustain the growth of auxotrophic strains of public or private good amino acids (Fig. 4A), using viable auxotrophs that could be generated (Fig. S4). The auxotrophic strains of public goods included *bat1Δ*/*bat2Δ* (auxotrophic for all branched chain amino acids), *leu2Δ*/*bat1Δ* (also auxotroph for all branched chain amino acids), *trp*5*Δ* (tryptophan auxotroph), while the private good auxotrophs included *met6Δ* (methionine), *lys1Δ* (lysine), *his3Δ* (histidine) auxotrophs (Fig. S4). To experimentally test this, we grew wild-type cells in synthetic minimal medium for 24 hours, and collected the culture supernatant (Fig. 4A). To this culture supernatant, we added only glucose and ammonium salts to make conditioned supernatant, and then assessed the growth of different auxotrophic strains in conditioned supernatant (Fig. 4A). For the public good auxotrophic strains, we assessed the growth of *bat1Δ*/*bat2Δ*, *leu2Δ*/*bat1Δ*, and *trp5Δ* cells in conditioned spent medium, including positive controls where the respective auxotrophic amino acids were supplemented at 2mM (Fig. 4B). Notably, the conditioned supernatant supported the growth of these public good auxotrophic strains, at times as effectively as when these cells were grown in conditioned supernatant supplemented with excess of the specific amino acid (Fig. 4B). These findings reveal that the public good amino acids are produced/secreted in sufficient amounts to satisfy the demands and support the effective growth of their respective auxotrophic cells (Fig. 4B). This further suggests that these amino acids are made by cells in amounts exceeding their own demands. We next assessed the growth of private good auxotrophs (*met6Δ*, *lys1Δ*, *his3*Δ) in conditioned supernatant. In contrast to the public good amino acid auxotrophs, the private good amino acid auxotrophic strains showed negligible growth in conditioned supernatant, in the absence of exogenously supplementing these respective amino acids (Fig. 4C). This demonstrates that *met6Δ*, *lys1Δ*, and *his3Δ* cannot obtain sufficient amounts of these respective amino acids from the spent medium to sustain growth (Fig. 4C).

**Fig. 4:**
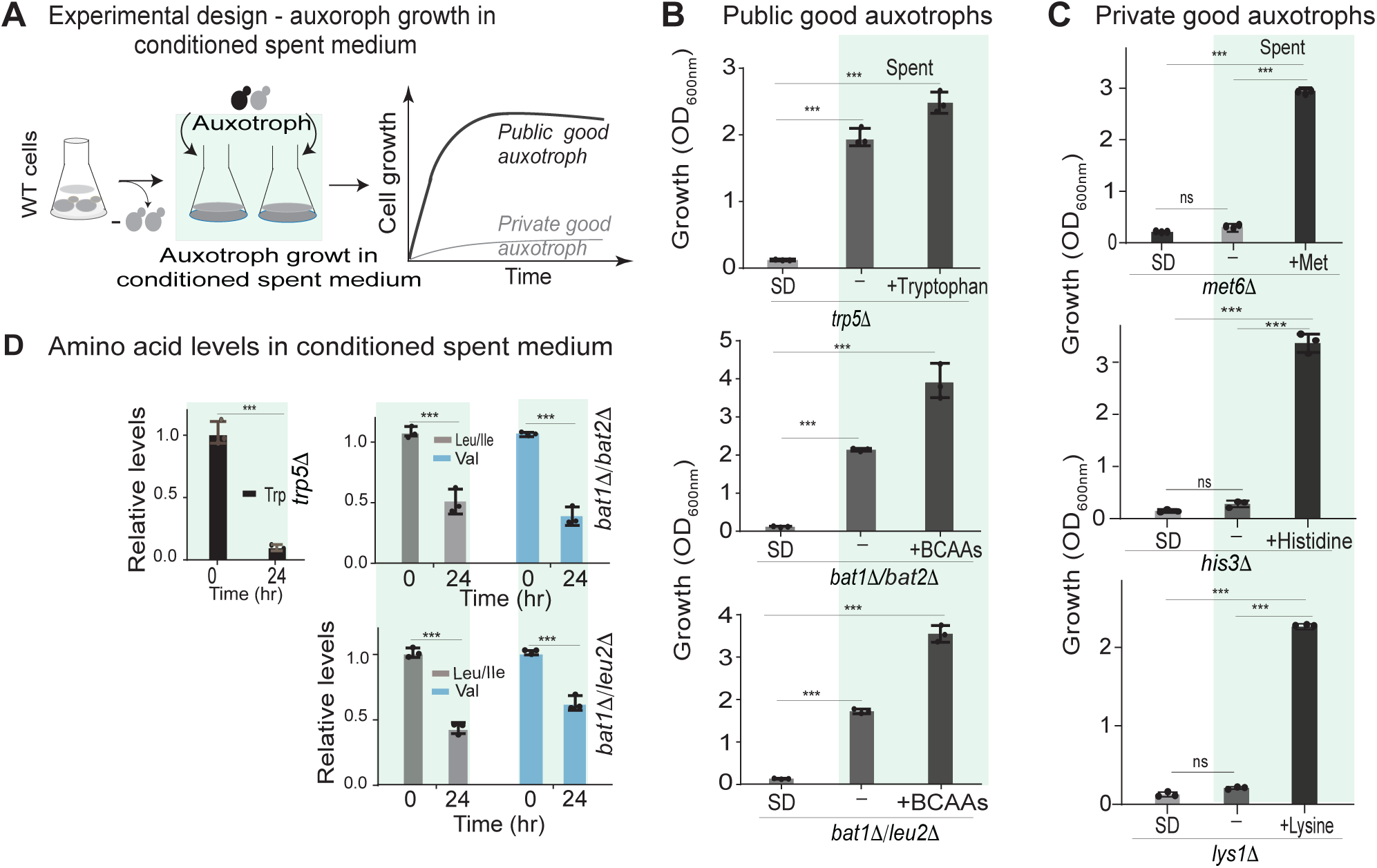
Cells secrete sufficient excess of amino acids to support growth of public good auxotrophs. A. Experimental design to assess auxotrophic growth in spent medium. Spent medium after 24hr growth of wild-type cells was collected and supplemented with glucose and ammonium salts to make conditioned supernatant. Subsequently, the specified public good auxotroph or private goods auxotroph is allowed to grow in conditioned supernatant. The predicted growth of a public good amino acid auxotrophic cells versus a private public good amino acid auxotrophic cells in conditioned supernatant is also illustrated. Also see Fig. S4 for auxotrophic cell growth. B. Growth of public good auxotrophs in spent medium: Auxotrophs of the following public good amino acids - *trp*5Δ (auxotrophic for tryptophan), *Δbat1*Δ*/Δbat2*Δ (auxotrophic for all branched chain amino acids), *bat*1Δ/*leu*2Δ (alternate of auxotrophic for BCAAs) – were grown in conditioned supernatant. As controls, these cells were also grown in conditioned supernatant that was supplemented with 2mM of the amino acid relevant to the specific auxotrophy. C. Growth of private good auxotrophs in conditioned supernatant: Auxotrophs of the following public good amino acids *met6*Δ (auxotrophic for methionine), *his3*Δ (auxotrophic for histidine), *lys1*Δ (auxotrophic for lysine) – were grown in spent medium supplemented with glucose and ammonium salts only. As controls, these cells were also grown in conditioned medium that was supplemented with 2mM of the amino acid relevant to the specific auxotrophy. D. Relative amounts of the specific extracellular amino acid (relevant to the auxotrophy) after 24hr of growth in conditioned supernatant. Auxotrophic cells were grown in conditioned supernatant (as in panel B). The bar plots show the relative amounts of the indicated amino acid in the extracellular environment, after the specified amino acid auxotroph was grown in conditioned supernatant. The auxotrophic strains used were: Δ*trp*5, Δ*bat*1Δ/bat2Δ, *leu*2Δ/*bat*1Δ. Each auxotroph was grown in spent medium of wild-type cells, and the amount of the specific amino acid was measured at 0hr and after 24hr. Data in all panels are from three independent biological replicates, represented as means ± SD, n = 3. *p<0.05, ***p<0.0001 (Student’s t-test).

Finally, to determine the actual utilization of these public amino acids by their auxotrophs, we assessed the relative levels of specific amino acids (tryptophan, leucine/isoleucine, and valine) in conditioned supernatant, after the growth of the indicated public good auxotrophic strain in this medium (Fig. 4D). After 24 hours of growth of the respective auxotrophic strain in conditioned supernatant, tryptophan, leucine/isoleucine, and valine levels significantly reduced (Fig. 4D). This indicated that substantial amounts of these amino acids are consumed from the extracellular medium by the respective auxotrophic strains, and reiterates that these amino acids were produced in sufficient amounts by wild-type cells to satisfy the demand requirements of auxotrophic cells (Fig. 4D). Collectively, these findings reveal that auxotrophic yeast cells can effectively consume/utilize only public good amino acids from the spent media, to meet their growth requirements.

### Growth of synthetic coculture communities of public-public and public-private good auxotrophs

Exchange based auxotrophic communities can form in yeast (6, 11, 13), yet underlying metabolic rules driving these interactions are unresolved. In particular, can pairs of auxotrophs of surplus public goods establish effective communities, based on the nature of the auxotrophy and the type of amino acid exchanged? Will auxotrophs of public good amino acids therefore form effective, mutual exchange-based communities when paired? Our data indicate that public good auxotrophs grow effectively by using the corresponding amino acid obtained solely from spent medium of prototrophic cells. We therefore asked if effective synthetic communities based on complementary amino acid exchange could be created, using paired co-cultures of public good-public good, or public good-private good auxotrophic strains. To address this, we curated pairs of public good auxotrophs, as well as pairs of public good-private good auxotrophs (Fig. 5A). We then inoculated each paired combination, with 50:50 ratios of each strain in low cell density in conditioned supernatant, and allowed these cells to grow together in this medium. The synthetic communities of auxotrophic-pairs used were: public good -public good auxotrophs (*bat1Δ*/*bat2Δ* + *trp*5Δ, and *bat1*Δ/*leu*2Δ + *trp*5Δ), and public good-private good auxotrophs (*bat*1/*bat*2Δ + *met*6Δ, and *bat*1Δ/*leu*2Δ + *lys*1Δ) (Fig. 5A). We monitored the fractional composition (of each auxotroph) over time (Fig. 5B and C). In order to accurately estimate the fraction of each auxotroph, each strain was engineered with a constitutive, fluorescent protein-label (mCherry or mNeonGreen) for all the public and private good auxotrophic strains, in the indicated combinations (Fig. 5B and C). For all public good-public good auxotrophic pairs of co-cultures, we observed robust growth and a stable population of each strain (Fig. 5B). For each of these pairs tested, a constant ratio of each genotype was maintained over 48 hours (Fig. 5B). In contrast, for public good-private good auxotrophic co-culture pairs, the fractional composition of each strain rapidly changed (Fig. 5C). The public good auxotrophic strains utilized the conditioned supernatant and grew, increasing in fractional composition, while private auxotrophic strains failed to thrive in the co-culture (Fig. 5C), with their relative population abundance decreasing to <5% (Fig. 5C).

**Fig 5.**
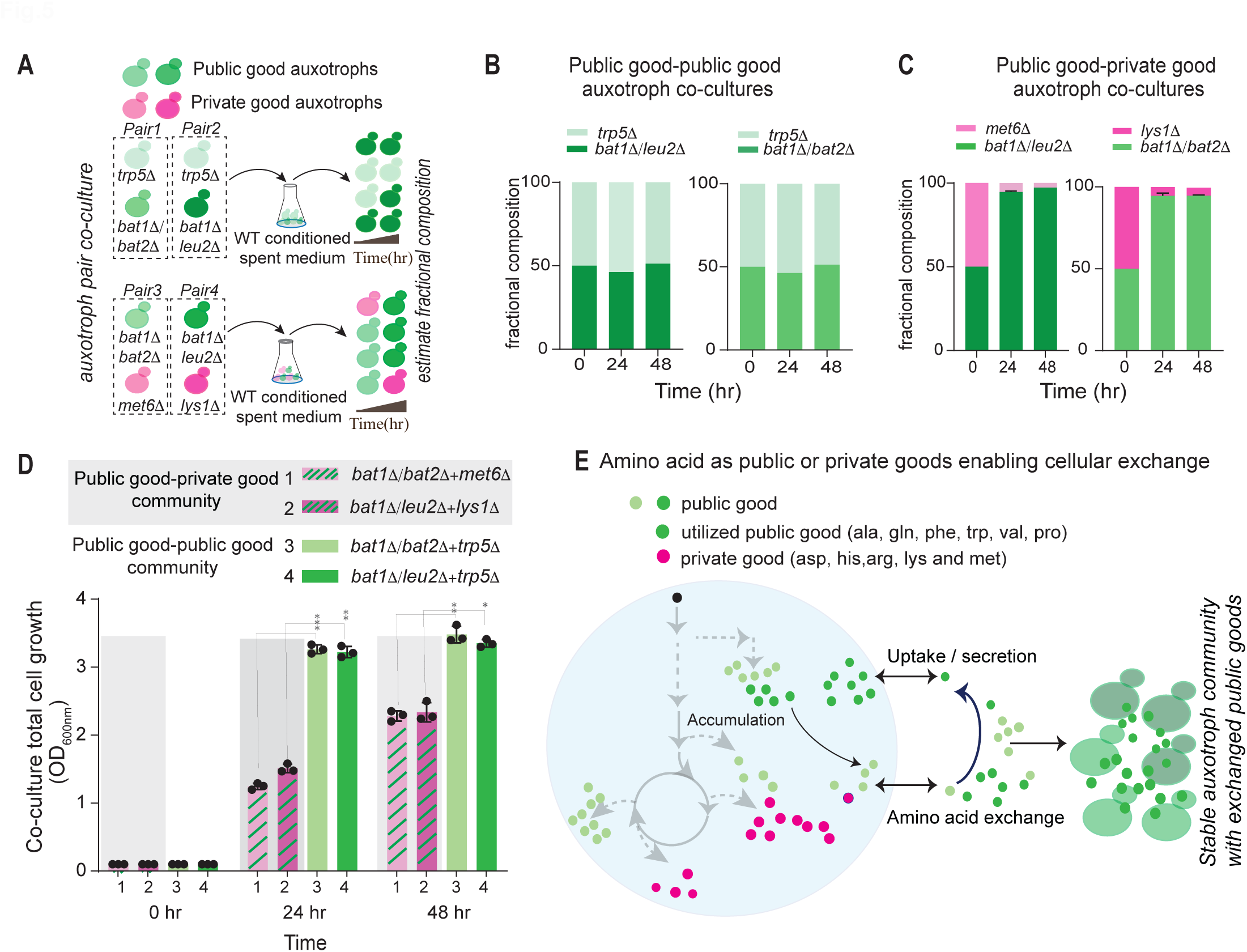
Synthetic communities of public good-public good auxotrophic pairs show synergistic growth based on mutual exchange. A. Experimental design to assess the growth of auxotroph pairs of co-cultures seeded in conditioned supernatant. The indicated combinations of public good-public good auxotroph pairs, or public good-private good auxotroph pairs were co-cultured in conditioned supernatant. The fractional (proportional) composition of each strain in the paired culture, as well as the total biomass attained, after the indicated time of growth is then estimated. B. Public good-public good auxotroph pairs maintain proportional growth: Bar plots displaying the fractional composition of each strain in the indicated paired co-cultures of public good-public good auxotrophs. The paired auxotrophic cell co-cultures tested were *trp5*Δ + *bat1*Δ*/leu2*Δ, and *trp5*Δ+*bat1*Δ*/bat2*Δ. C. Private good auxotrophs are out competed by public good auxotrophs: Bar plots displaying the fractional composition of each genotype in paired co-cultures of public good-private good auxotrophs. The paired auxotrophic cell co-cultures were *met6*Δ + *bat1*Δ*/leu2*Δ, and *lys1*Δ + *bat1*Δ*/bat2*Δ. D. Public good-public good auxotrophic pairs show synergistic growth: Total biomass at the indicated time after growth of public good-public good auxotrophic pairs, compared to public good-private good auxotrophic pairs. Each paired community was grown in the same conditioned supernatant. The public good-private good auxotrophic pair communities tested were *bat1*Δ*/bat2*Δ + *met6*Δ and *bat1*Δ*/leu2*Δ + *lys1*Δ. The public good-public good auxotrophic pair communities tested were *bat1*Δ*/bat2*Δ + *trp5*Δ and *bat1*Δ*/leu2*Δ + *trp5*Δ. E. A schematic model illustrating how private good or public good amino acids function to shape exchange-based communities: Amino acid production, secretion, and accumulation and exchange in *S. cerevisiae* are illustrated, and amino acids can be clearly identified private goods (pink balls) or public (green balls) goods. The auxotrophs of public good amino acids that are produced and secreted in excess are effective in forming successful, exchange based communities, that show enhanced growth compared to single public-good auxotrophs or those paired with private good auxotrophs. Data in all experiments are from three independent biological replicates (n=3), represented as mean ± SD. *p<0.05, ***p<0.0001 (student’s t-test).

We next asked if pairing public good-public good auxotrophs in co-cultures led to a gain in cell numbers and overall growth that potentially comes from amino acid exchange (Fig. 5). We also asked if the growth can be explained by the conditioned supernatant alone, or indicated reliance on secreted amino acids (Fig. 5D). For this, we estimated cell numbers of the respective paired co-culture over time (Fig. 5D). We observed that the public good-public good auxotrophic pairs grew robustly, reaching significantly higher total biomass in 24 and 48 hours of co-culture (Fig. 5D). These data indicate that co-operative amino acid exchange between the public good-public good auxotrophs leads to a significant increase in population size and biomass, unlike with public-private co-cultures that rely exclusively on available amino acids in conditioned supernatant (Fig. 5E). The public good-public good auxotroph co-cultures easily outperform public-private co-cultures in biomass formation, through a robust exchange of public goods (Fig. 5E).

Collectively, these data underscore the importance of identifying public good amino acids and their auxotrophs, through which exchange based communities can be established (Fig. 5E). Public good amino acids, produced in excess, are highly effective in supporting the growth of their auxotrophs, and can be used by cells to form effective, exchange based synthetic communities (Fig. 5E).

## Discussion

In this study, we elucidate the quantitative amounts, and temporal dynamics of the formation, secretion and consumption of amino acids during yeast cell growth. We identified phases of high amino acid biosynthesis, which are restricted to the early phase of cell growth in batch culture, and gene expression programs induced at later time points will reflect a starvation response by cells attempting to restore biosynthetic flux without being able to do so (Fig 1D). Subsequently, we built a quantitative blueprint of intracellular amino acid amounts and dynamics, identifying that the relative proportions of amino acids change substantially over different phases of growth (Fig 2C). Interestingly, the extracellular environment was entirely distinct from the intracellular pools (Fig. 2F). Through this, we identified the amino acids that are tightly privatized by cells, versus those that are publicly exchanged. Interestingly, the privatized amino acids are required for continuous, distinct metabolic and growth needs in cells (Fig. 3C). The public good amino acids themselves form two groups – those that are produced and secreted in surplus, and those that are produced in surplus initially but are later consumed (Fig. 3B). From these data, we find that auxotrophs of public goods can grow effectively using resources exclusively obtained from their extracellular environment (Fig. 4B). These public good auxotrophs are also ideally suited to form stable, exchange-based synthetic communities (Fig. 5D). Collectively, we can now build a quantitative blueprint of the hierarchies in a cellular amino acid economy, built on biosynthesis, secretion and consumption (Fig. 5E). The highlight is direct evidence for distinct prioritizations of privatized amino acids, based on continuous cellular needs, as well as a truly dynamic intracellular amino acid economy that has been largely underappreciated (Fig. 5E).

Even in versatile laboratory models or industrially important cells like yeast or *E. coli*, our understanding of the intracellular and extracellular amino acid economy is sparse. Current knowledge is limited to static snapshots of certain intracellular or extracellular amino acids or other metabolites (23, 30–32). It is clear that cells have dramatically different amounts of individual amino acids (23), or maintain distinct amino acid reserves (24). Further, the amino acid economy is shaped not by the metabolic cost of an individual amino acid, but instead by total demand for that amino acid coupled with the ease of synthesis (22). This makes the amino acid economy especially dynamic and constantly changing in a growing cell. The extent and nature of this dynamic cellular economy is a feature of both the inherent constraints imposed by growth, as well as the metabolic flexibility of that cell. In light of our findings, made with reductionist approaches in yeast, it is interesting to ask what the range and limits could be for amino acid amounts inside different types of cells, and in what contexts would changes in one amino acid result in the induction or reduction of another amino acid.

Cooperative interactions based on syntrophic metabolic exchange and cross feeding enable groups of cells to build metabolic efficiencies, drive auxotrophic cell populations towards new metabolic imbalances and interactions, or even improve lifespan and survival (3, 9, 10, 31, 33). Designing and engineering synthetic communities based on syntrophic exchange is an area of considerable interest, for the obvious potential it holds for biotechnological applications. Yet many efforts to design exchange based microbial consortia for biotransformations fall short of meeting expectations (12, 34, 35), in part due to challenges in quantitatively untangling metabolic interactions (3). In yeast, combinatorial efforts, or selection and adaptation, have been used to build exchange based communities (6, 11, 13, 33). In contrast, our study offers a bottom up approach towards building effective microbial consortia based on metabolic exchange, which would require a quantitative understanding of the metabolite economy – based the production, secretion or consumption of this metabolite, or its intermediate. This is critical in the context of the cellular amino acid economy, where identifying supply versus demand constraints for distinct amino acids (22) will define metabolic frameworks that determine which amino acids or their intermediates might be effectively exchanged, or can build exchange-based communities. Context specific, quantitative insights into their production, secretion, consumption cycles can therefore identify resource-bottlenecks, and suggest ways to overcome these bottlenecks. This quantitative approach towards defining the amino acid economy will broaden our understanding of resource allocation strategies within and between cells, and can inform the rational design of metabolically engineered cell consortia as factories.

## Materials and Methods

### Yeast growth and media conditions

A prototrophic, haploid (CEN.PK mat a) strain of *Saccharomyces cerevisiae* (36) was used in all the experiments, and was maintained in YPD medium containing yeast extract (1%), peptone (2%), glucose (2%) and agar (2%). For growth experiments, a primary culture of *S. cerevisiae* was grown overnight in YPD nutrient broth in a screw-capped tube and the primary culture was washed and diluted to an OD_600nm_ ∼0.10/0.20 of cells, in the growth medium. Synthetic defined medium (SD) used contained nitrogen base without amino acids and 2% glucose (filter sterilized), with glucose as the sole carbon source and ammonium sulfate (as per media LoT-7109979) as the sole nitrogen source. Cells were incubated in an orbital shaker operating at 240 rpm at 30°C. The list of strains and plasmid used in this study are provided in Supplementary Table S1 and S2.

### Intracellular amino acid extraction and detection

Intracellular amino acids were extracted and quantitatively estimated using liquid chromatography– tandem mass spectrometry (LC-MS/MS) approaches described earlier (37), with samples collected from cultures grown in synthetic defined minimal medium over a 24 hours growth period. Specifically, at time points of (2hr, 4hr, 8hr, 12hr, and 24hr) equal numbers of cells (∼2×10^7) were quenched in extraction buffer (60% methanol), extracted in 75% ethanol and dried down using speed vacuum (rotatory evaporator). Metabolites were dissolved in 300µl mass spectrometry grade water and 10µl sample was injected for LC-MS/MS and separated using Synergi 4-µm Fusion-RP 80 Å (150 × 4.6 mm) LC column (Phenomenex, 00F-4424-E0). Solvents used for amino acid and TCA derivatives are 0.1% formic acid in water (Solvent A) and 0.1% formic acid in methanol (Solvent B). The LC-MS/MS (mass spectrometer) used was an AB Sciex QTRAP 5500 with Shimadzu Nexera series UPLC system. Detection of amino acids was done in positive polarity mode. Mass spectrometry data were acquired using analyst 1.6.2 software (Sciex). For analysis, Multi-Quant version 3.0.1 and Peak View version 2.0 were used. A table with Q1 and Q3 parameters for all metabolites including labelled and unlabelled forms is provided (supplementary Table S3). To calculate intracellular amino acids per cell, the following information was considered 1.0 ml of 1.0 OD_600nm_ culture corresponded to ∼2×10^7 cells, and the volume of a single yeast cell ∼40×10^−15^ fl was used. The concentration of intracellular amino acids (µM) = 1 x (cells conc. in µM/L) / (Molecular weight of individual amino acids/AA) x 2 x (10^7 x 40 x 10^−15^ L)

### Extracellular amino acid extraction and quantification

*Saccharomyces cerevisiae* cultures were grown in a minimal medium from early to the post-diauxic growth phase, as described earlier. At specific time points (2, 4, 8, 12, and 24 hour), 1.0 ml culture was withdrawn. The culture was centrifuged for 5 minutes at 7000 rpm at RT and 250/500µl supernatant was used for extracellular metabolite extraction. The extracellular amino acids were extracted with 75% ethanol. The extracted metabolites were then vortexed for 1.0 minutes and placed on ice for few minutes. The samples were centrifuged at 16000 rpm for 12 min at RT and 750 µl of the supernatant was collected in new eppendorf tubes, which were dried using a speed vacuum. The dried metabolites were stored at –80°C until mass spectrometry analysis was performed as described earlier (37).

### Auxotrophic strain generation

In the prototrophic background, we introduced multiple knockouts through homologous recombination of specific genes using standard drug selection cassettes (38). Correct clones were verified using PCR with specific primers. The list of auxotrophic strains used in this study are provided in see supplementary Table S2.

### 13C4-aspartate utilization flux measurement by LC-MS/MS

The ^13^C_4_-aspartate utilization flux was measured by tandem mass spectrometry to investigate its incorporation into intermediates of the TCA cycle, gluconeogenesis and other metabolites. Cells were grown in a minimal medium (SD; synthetic defined media) for 4hr and 24 hours, pulsed with 1mM ^13^C_4_-aspartate (Cambridge Isotope Laboratories, CLM-1801-0.25G) and incubated for 20 minutes. Intracellular metabolites were extracted from ∼1.0 OD cells and derivatized as reported earlier (37). Samples were dissolved in 1.0 ml methanol and water (ratio 1:1) and a 10µl sample was injected for HPLC separation using methods described earlier (37). Detection of label incorporation into TCA intermediates was done in positive polarity mode. Parent (Q1) and product ions (Q3) that were used for detection of labelled metabolites are provided in supplementary Table S3. The total ^13^C_4_-aspartate label incorporation was calculated as the sum of all individual peak areas coming from ^13^C_4_-aspartate incorporated and detected for each specific metabolite. Relative label incorporation was calculated by normalizing with respective control values in each set.

### Measurement of deuterated Alanine (d4-alanine) conversion to pyruvate by LC-MS/MS

Deuterated alanine (d4-alanine) uptake and incorporation into pyruvate was measured using tandem mass spectrometry. The culture was initially grown in minimal medium, at 4hr and 24 hours before being pulsed with 1mM d4-alanine (Cambridge Isotope Laboratories, 299286-20G) and incubated for 5 minutes. After that, metabolites were extracted as described earlier (37), and d4-labelled pyruvate (coming from d4-alanine) was estimated, similar to as described above. Detection of d4-pyruvate used the Q1 and Q3 masses described in supplementary Table S3. Statistical significance was calculated using an unpaired Student’s t-test.

### Luciferase reporters for Gcn4 activity

To measure Gcn4 activity, we used a well-established Gcn4-luciferase reporter system (Supplementary information Table S1) as described earlier (22, 39). Cells expressing the reporter were grown in the presence of antibiotic (G418), and the relative luciferase expression was estimated in cells grown at the respective minimal medium and collected at the indicated time. For the experiment, overnight cultures with cells expressing the reporter were collected and washed twice in minimal media, and used to set up a fresh culture in 250 ml flask, with 100 ml synthetic defined medium with 2% glucose. One groups of flasks served as a control with 2mM amino acid (+AA) supplemented. In the control groups, after 2 hours, 2mM amino acid were added and after 4hr, cells were collected in at an OD_600nm_ ∼10. Biological replicates of samples at 2, 4, 8, 12, and 24 hours were collected. Collected cell pellets were washed with 300μl lysis buffer containing 1xPBS (137 mM NaCl, 2.7 mM KCl, 10 mM Na2HPO4, and 1.8 mM KH2PO4 and 1 mM PMSF, pH 7.4) twice and stored at −80°C till the luciferase assay was performed. Pellets were re-suspended in 300μl lysis buffer, proteins were extracted by bead-beating with lysates maintained on ice, and extracted protein concentrations were estimated by BCA (bicinchoninic acid) protein assay kit. Equal total protein concentrations of lysates were used while measuring the luciferase activity, using 2μg/μl protein for 20μl reaction. Luciferase activity was measured through a luciferase assay kit (Promega, E1500) and a luminometer (Sirius, Titertek Berthold Detection systems). The luciferase activities for each replicate (Relative Light Units per Sec (RLU/sec)) were normalized with individual controls (+AA). The relative RLU in luciferase activities between different time points in minimal media were used to estimate changes in the active translation of Gcn4 transcription factor proteins.

### Growth assays

The growth of auxotrophic strains was monitored in synthetic minimal medium (as described earlier) agar plates. Primary cultures of auxotrophic cells were washed, resuspended in 1 ml medium at an OD_600nm_ of 1.0, serially diluted 10X into multiple tubes, and a 5μl drop from each 10X dilution was spotted onto minimal medium agar plates without amino acids, or with the respective (auxotroph) amino acid supplemented. Auxotrophic strains used are listed in Supplementary Table S2.

### Assay for growth using conditioned supernatant

Prototrophic wild type *S. cerevisiae* cells were grown in minimal medium with 2% glucose, without amino acids for 24 hours at 30 °C in shaker incubator. The supernatant was harvested after 24 hours of growth, which was filtered using a 0.22μm membrane. To this spent medium, only glucose (2%) and ammonium sulfate (∼35mM) were added to make ‘conditioned supernatant’. This was used as the media source to test the growth of the respective auxotrophic strains (Supplementary Table S2). For growth experiments, a primary culture of the specified auxotrophs were grown overnight in YPD nutrient broth, and the primary culture was collected, washed and diluted to an OD_600nm_ ∼ 0.10, and grown in the conditioned supernatant at 30 °C in the shaker incubator.

### Co-cultures of private-public and public-public auxotroph goods

Conditioned supernatant (described above) was used for individual strain culture, as well as for co-cultures of combinations of different auxotrophic cells. Combinations of auxotrophic cell pairs were seeded at an OD_600nm_ of 0.05 each in the conditioned supernatant. In co-cultures, the starting ratios of the two auxotrophs was 50:50 (ratio), and at a starting OD_600nm_ of 0.1, grown in 100 ml flasks containing 20 ml conditioned supernatant. The growth of the co-cultures, as well as their fractional compositions were determined by counting colony forming units (CFU), as well as counting cells with reporter expression, where cells were labeled with green (mNeonGreen) or red (mCherry) fluorescent tags at an endogenous locus (Supplementary Table S2).

### Cell counting

Co-cultured cells expressing a fluorescence reporter in conditioned supernatant were visualized using a microscope (Olympus BX53) with a 20 X air objective. The fraction of cells expressing each reporter was then estimated by counting.

### Data representation and statistical analysis

Graphs were plotted and data analyzed using GraphPad Prism 6. Two-tailed Student’s *t* test was used to estimate statistical significance unless otherwise specified. For the flux experiments, the raw intensity values for each replicate were normalized to respective controls. For the mass spectrometry experiment examining relative levels in spent media and conditions spent media was used to estimate statistical significance. P values and *n* for corresponding experiments have been specified in figure legends.

## Author Contributions

S.A., and S.L, conceived and designed the project. S.A performed the experiments. S.A., S.L, discussed and analyzed the data. S.A., S.L, drafted the manuscript. S.A., S.L, edited the manuscript. All authors have read and approved the manuscript.

## Acknowledgements

We acknowledge extensive use of the inStem /NCBS/ CCAMP mass spectrometry facilities for HPLC and mass spectrometry. We thank Deepa Agashe, Aswin Seshasayee, Anjana Prasad, Christian Kost and members of the SL Lab for critical discussions and comments on the manuscript. SA acknowledges DST-SERB N-PDF File Number: PDF/2022/000700. SL acknowledges a DBT-Wellcome Trust India Alliance Senior fellowship (IA/S/21/2/505922), the DBT S. Ramachandran National Bioscience Award for Career Development, and the DBT-DFG Indo-German collaboration grant (IC-12025(22)/4/2023-ICD-DBT) from the Department of Biotechnology, Govt. of India for support.

